# Effects of Zika Virus Non-Structural Proteins on the Innate Antiviral Response

**DOI:** 10.64898/2026.11.21.765188

**Authors:** Santiago Ochoa Zermeño, Ava C. Lariego, Rori Hoover, Christopher L. Plaisier, Benjamin B. Bartelle

## Abstract

Innate immune pathways are therapeutic targets of prime importance, both for antiviral and neurodegenerative therapies. The Zika virus is highly evolved to manipulate these same pathways so expertly that half of infections go unnoticed. While the direct inhibition of immune effectors by ZIKV nonstructural proteins is well characterized, second order effects on immune responsive cellular functions are less understood. We induced the innate antiviral response in human microglia like cells, perturbed that response using 6 ZIKV nonstructural genes, and resolve systems level effects using scRNA-seq and hdWGCNA. We recapitulate known ZIKV bioactivity and identify specific suppression of the PA28 immunoproteasome by NS2B, demonstrating how ZIKV can shape tissue level neuroinflammation.

## Introduction

Viral infections cause neuroimmune activation via mechanisms that have also been identified as chronic drivers of neurodegeneration. (Salter & Stevens, 2017; Hammond et al., 2019; Paolicelli et al., 2022). Most fundamentally, the RIG-I-like receptor (RLR) and mitochondrial antiviral signaling (MAVS) signalosome responding to cytosolic double stranded (ds) dsRNA, and the cGAS-STING complex responding to cytosolic dsDNA. Both pathways free NF- κB or activate interferon regulatory factor 3 (IRF3). Chronic activation of cGAS-STING is a known driver of Alzheimer’s disease (Zhang et al. 2025). TANK binding kinase (TBK1) activation, recruited by MAVS, is coincident with Amyotrophic Lateral Sclerosis (ALS) and Frontotemporal Dementia (FTD) (Ahmad et al. 2016). Parkinson’s-associated proteins like PINK1 interact with IRF3 to modulate RLR activation to accelerate disease progression (Fan et al. 2025). This overlap between viral response and neurodegenerative mechanisms makes antiviral pathways valuable targets for the treatment and prevention of neurodegenerative disease.

Conversely, viruses have evolved specifically to disarm innate immunity, with HIV famously avoiding any double stranded forms outside of the nucleus. Zika virus (ZIKV), while best known for infecting neural progenitor cells (hNPCs), is both neuro and immunotropic. (Tang et al., 2016; Retallack et al., 2016; Dello Russo et al., 2018). ZIKV efficiently infects and replicates in the brain’s dedicated innate immune cell with antiviral sensing and response mechanisms well beyond other cell types (Lawrence et al. 2019). Unlike lentiviral HIV, flaviviral ZIKV must form dsRNA as part of its lifecycle, thus should strongly activate RLRs. Instead, ZIKV encodes multifunctional non-structural (NS) proteins that selectively target and disassemble innate immune cascades (Barrows et al., 2018; Serman & Gack, 2019; Lee et al., 2020). ZIKV NS2A and NS4A strongly suppress MDA5-, RIG-I-, MAVS-, TBK1-, and IRF3-mediated activation of NF-κB and IFN-β promoter activity (Nguyen et al., 2019; Lee et al., 2020). The ZIKV NS2B-NS3 viral protease complex directly targets and degrades key adaptor proteins, including MAVS, Jak1, and STING/MITA, thereby inhibiting downstream IRF3 phosphorylation and type I interferon induction (Li et al., 2019; Serman & Gack, 2019; Lee et al., 2020). Additionally, ZIKV NS1 suppresses host interferon induction, while NS4A and NS4B co-operatively dysregulate host Akt-mTOR signaling pathways to suppress cellular stress responses and promote viral survival (Liang et al., 2016; Xia et al., 2018; Lee et al., 2020).

ZIKV tropism for microglia is due to specificity for integrin αvβ5 and the TAM receptor AXL via Gas6 cytokine bridging (Nowakowski et al., 2016; Meertens et al., 2017; Wang et al., 2020). Gas6/AXL is upregulated in microglia by the Type I interferon response and its signaling is innately anti-inflammatory (Meertens et al. 2017). ZIKV cloaking in Gas6 is only the first step in a chain of immunomodulatory mechanisms ZIKV leverages in order to subjugate the brain’s dedicated antiviral cell type

Given the diversity of immune evasion mechanisms already described, we took a systems view of ZIKV effects on innate immune signaling to put described mechanisms into context and to bioprospect for novel immunosuppressive ZIKV mechanisms. Here we took a modified Perturb-seq approach, dissecting the innate immune effects of the individual ZIKV non-structural proteins (NS1–NS5) for a transcriptomic view of how each viral modulator perturbs intrinsic host immune networks (Lee et al., 2020).

We recapitulated the microglia antiviral response using polyI:C (PIC) to trigger the molecular recognition of double stranded RNA, activating TLR3 and MDA5/RIG with secondary Jak/STAT activation (He et al. 2021). We utilized the immortalized human microglial cell line HMC3, which only partially recapitulates microglial biology, but possesses the full enzymatic machinery for TLR3, MDA5/RIG, RLR, NF-κB, and type I interferon signaling, offering a standardized model for fine precision immune studies (Janabi et al., 1995; Dello Russo et al., 2018).

This systems view of ZIKV interactions within HMC3 cells recapitulated known immunomodulatory bioactivity, while cataloging the second order effects of specific interactions. In addition, we identified novel bioactivity in the modulation of the proteasome and MHC Class 1 antigen display, suggesting immunomodulatory effects that extend beyond the infected cell.

## RESULTS

### HMC3 cells recapitulate canonical antiviral responses upon PIC stimulation

In order to understand the effects that ZIKV might have on antiviral response we first confirmed that PIC could trigger classical antiviral responses by transcriptomics. In differential expression analysis, between PIC, LPS, and CTRL, samples treated with PIC showed upregulation of interferon stimulated gene families (e.g.IFIT, ISG) and pro-inflammatory markers (e.g CCL5) (**Fig 1a,b**). Gene ontology (GO) enrichment analysis of upregulated genes in the PIC treated sample found an enrichment of innate immune response and defense response to virus gene sets.

Transcriptional upregulation in response to LPS showed almost total overlap with that found in response to PIC. With all but one of 89 upregulated genes in the LPS condition also being upregulated in the PIC condition. (**Fig S1**). This bears out in GO enrichment analysis finding both conditions share terms for innate immune activation and defense response to virus and bacterium. PIC responses shared all LPS genes with 7 fold more genes being upregulated. GO terms unique to PIC response point towards upregulation of cell junction loss, chemotaxis, angiogenesis, and T cell proliferation and recruitment. Indicating that HMC3 cells retain a robust antiviral response to PIC activation beyond stimulus of a single pathway.

To further characterize HMC3-PIC response we performed unsupervised clustering of PCA embeddings. We identified 14 clusters, 2 of which are almost exclusively made up of PIC treated cells (cluster 4 and 6). And one cluster being almost completely depleted of PIC treated cells (cluster 0) (**Fig 1c**). The marker genes for cluster 4 and 6 overlap heavily with genes involved in antiviral response. While cluster 0 was marker genes were associated with cell division.

**Figure 4.1.**
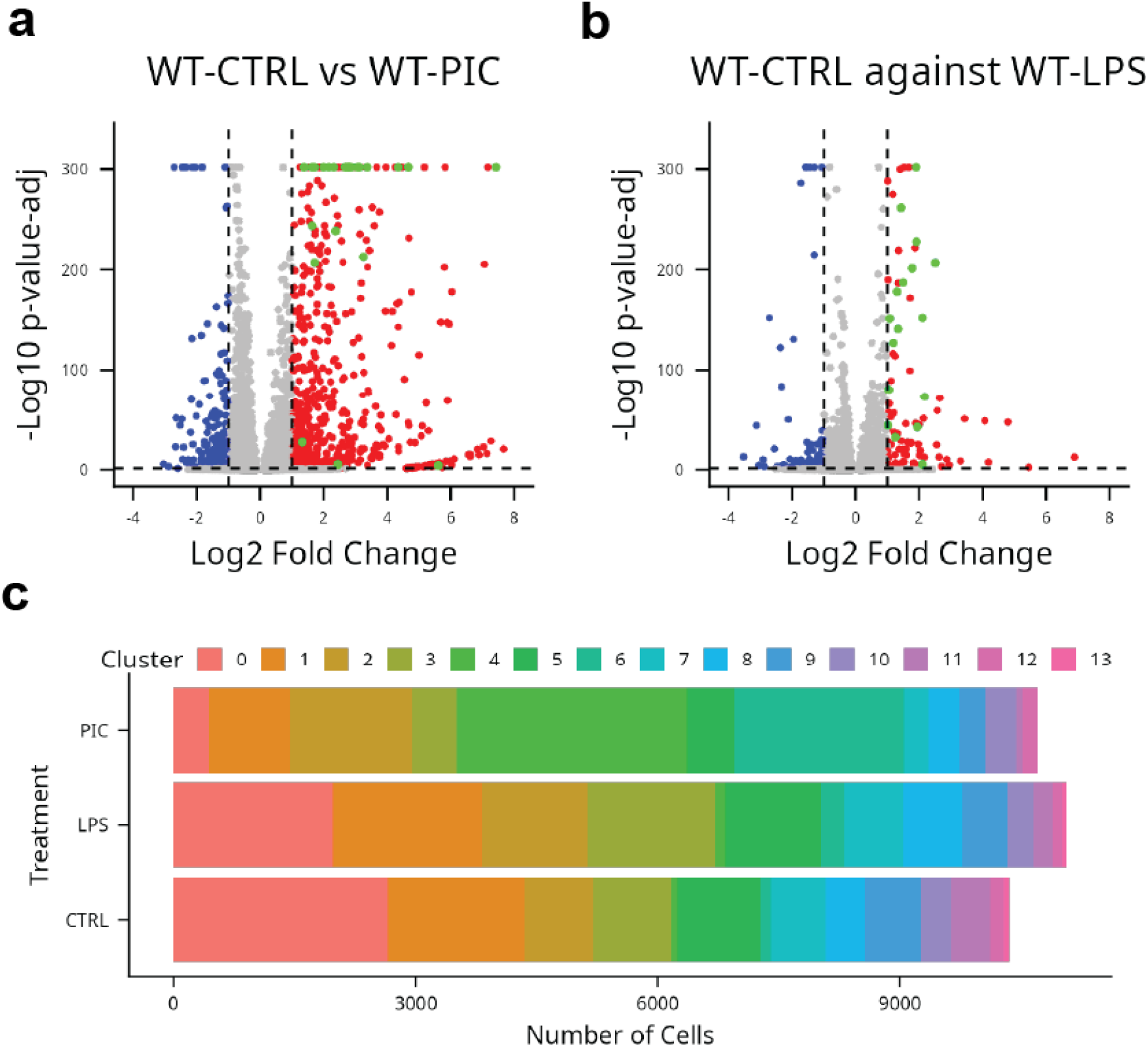
PIC and LPS produce distinct expression profiles. (a,b)Volcano plot showing differential gene expression of PIC and LPS treated WT cells against their isogenic control. Thresholds for significant differential expression were set at a log fold change >2, and p adjusted value <0.05, thresholds are depicted as dotted lines. Upregulated members of gene families involved in inflammation and cellular innate immunity (IFIT,ISG,CCL,OAS) are highlighted in green. (b) WT cells were subset and clustered using k nearest neighbor algorithm on PCA embeddings. Stacked bar plot shows number of cells belonging to each discovered cluster across treatments.

### The HMC3 antiviral response to PIC stimulation presents a highly correlated, transcriptional module

Having established a significant response to PIC stimulation in our cell line, we parsed the underlying functions of the antiviral response, using high dimensional Weighted Gene Correlation Network Analysis (hdWGCNA). We identified 9 modules in total across all conditions (**Fig 2a**). To determine which of these modules were capturing transcriptomic variance triggered by PIC response, we compared average module eigengene scores, as a measure of the common coexpression patterns of genes across sampled cells. Module eigengene scores decreased in the absence of their coordination by shared transcription factors. The turquoise module eigengene was only found in cells treated with PIC, while all the other modules are well represented across treatments. (**Fig 2 b**). Implicating the turquoise module as the sole module capturing coordinated gene expression involved in antiviral response. Hereafter the turquoise module is referred to as WT PIC module

Antiviral and inflammatory GO terms were highly enriched in this module. (**Fig 2c**). Other modules capture functions present across antiviral and homeostatic states, cell division, lipid metabolism etc. (**FigS2**). We compared the gene members of the WT PIC module against gene ontologies of defense response to virus (GO:0051607) and the differentially upregulated genes from a CTRL to PIC DE analysis to determine whether the WT PIC module captures different transcriptomic information from that measurable by differential expression (**Fig 2d**). The majority (66%) of genes differentially upregulated in the PIC condition versus CTRL, also appeared in the WT PIC module. 45% of genes annotated to the “defense response to virus” ontology term appear in the WT PIC module. In both cases the overlap with the WT PIC module is significant. A third of genes belonging to the WT PIC module do not overlap with either the DE gene set or to genes associated with the viral defense GO term.

**Figure 4.2.**
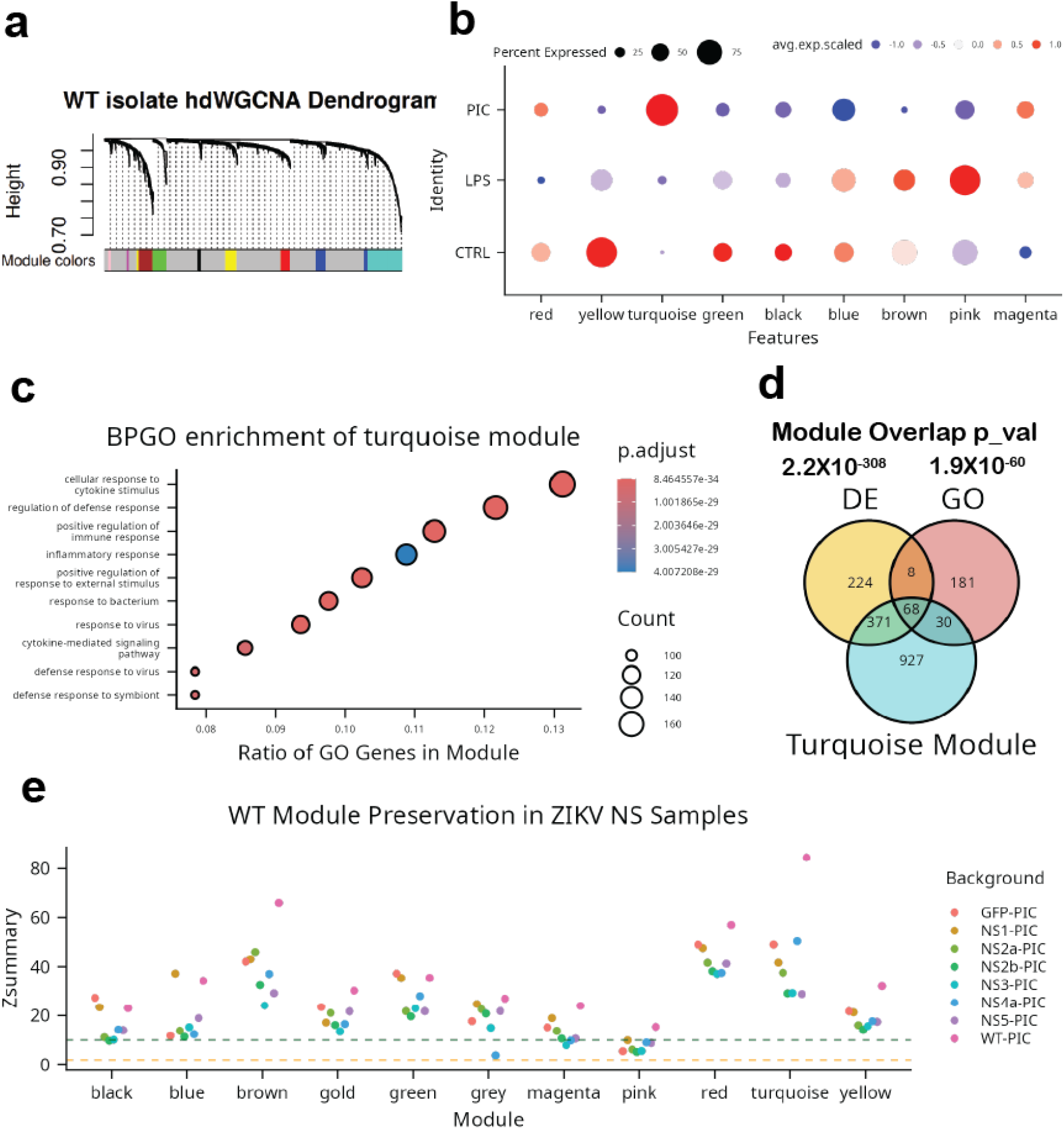
The turquoise module captures WT PIC response and is recapitulated in NS transduced samples. (a) hdWGCNA was performed on WT samples, producing 9 modules. Dendrogram branching depicts groups of genes with decreasing dissimilarity scores from modules of uniquely coexpressed genes.Lower bar shows module membership of genes in the dendrogram.(b)Dot plot showing module eigengene score in WT samples across all three treatments. Each row shows the average module eigengenge score of WT cells in that treatment group. Size of dot depicts percentage of cells in the group with a positive module eigengene score. The turquoise module eigengene is positively expressed disproportionately within the PIC treated sample. (c)Gene Ontology Enrichment Analysis was performed on the genes belonging to the turquoise module. The dot plot shows the ten enriched terms with the lowest adjusted p value, represented by the dot’s color. The amount of genes associated with the particular term is represented by the dot’s size. The x axis represents the ratio of genes associated to the GO term as a percent of the total genes belonging to the module (1396 genes)(d) Venn diagram displaying gene set overlap between genes upregulated in a DE analysis of WT-CTRL vs WT-PIC (yellow), genes belonging to GO term: 0051607 (red) and genes belonging to the turquoise module (turquoise). The number of overlapping genes is shown inside of the overlapping region. Significance of total overlap with the turquoise module is displayed over the name of the other two genesets.(e) Z scores of module preservation from WT hdWGCNA in all other backgrounds. Z scores <2 (dotted orange line) indicate modules not preserved in samples; while z scores >2 and <10 (dotted green line) indicate modules moderately preserved.

### The HMC3 antiviral transcriptomic response can be divided into 3 sub-modules

Having defined the wild type HMC3-PIC transcriptomic response we proceeded to determine how ZIKV proteins individually modulate it. Projecting both ZIKV NS1-5 transduced HMC3 cells with and without PIC stimulation into UMAP space did not result in novel clusters of cells with differential expression. All samples were well integrated across genetic backgrounds and showed considerable overlap based on treatment (**S3**). Despite known ZIKV NS interference in host antiviral activity, the WT PIC module was recapitulated in ZIKV NS1-5 transduced cells, however the module had increased variance, evidenced by a low module coherence score (**Fig 2e**).

We then subset data from HMC3 and ZIKV NS1-5 PIC and CTRL conditions, to focus only on transcriptional variance due to the antiviral responses and potential ZIKV modulation. This new approach produced 22 submodules, 3 of which were subdivided genes that formerly grouped to the WT PIC, representing concomitant functions underlying innate antiviral response (**Fig 3a**). The antiviral response submodules (seagreen, azure and orangered) were still unique to PIC stimulated cells and presented more specific GO enrichment and function specific hub genes:

The canonical antiviral submodule (seagreen) captured TLR3 signaling leading to interferon release and interferon stimulated gene expression (**Fig 3b**). The proteasome submodule (azure) captured transcription of genes involved in proteasome activation, key among them PSME1, and PSME2, and ribosome biogenesis (**Fig 3c**). The EGFR module (orangered) represents a growth factor response to EGF signaling via MAPK (**Fig 3d**).

**Figure 4.3.**
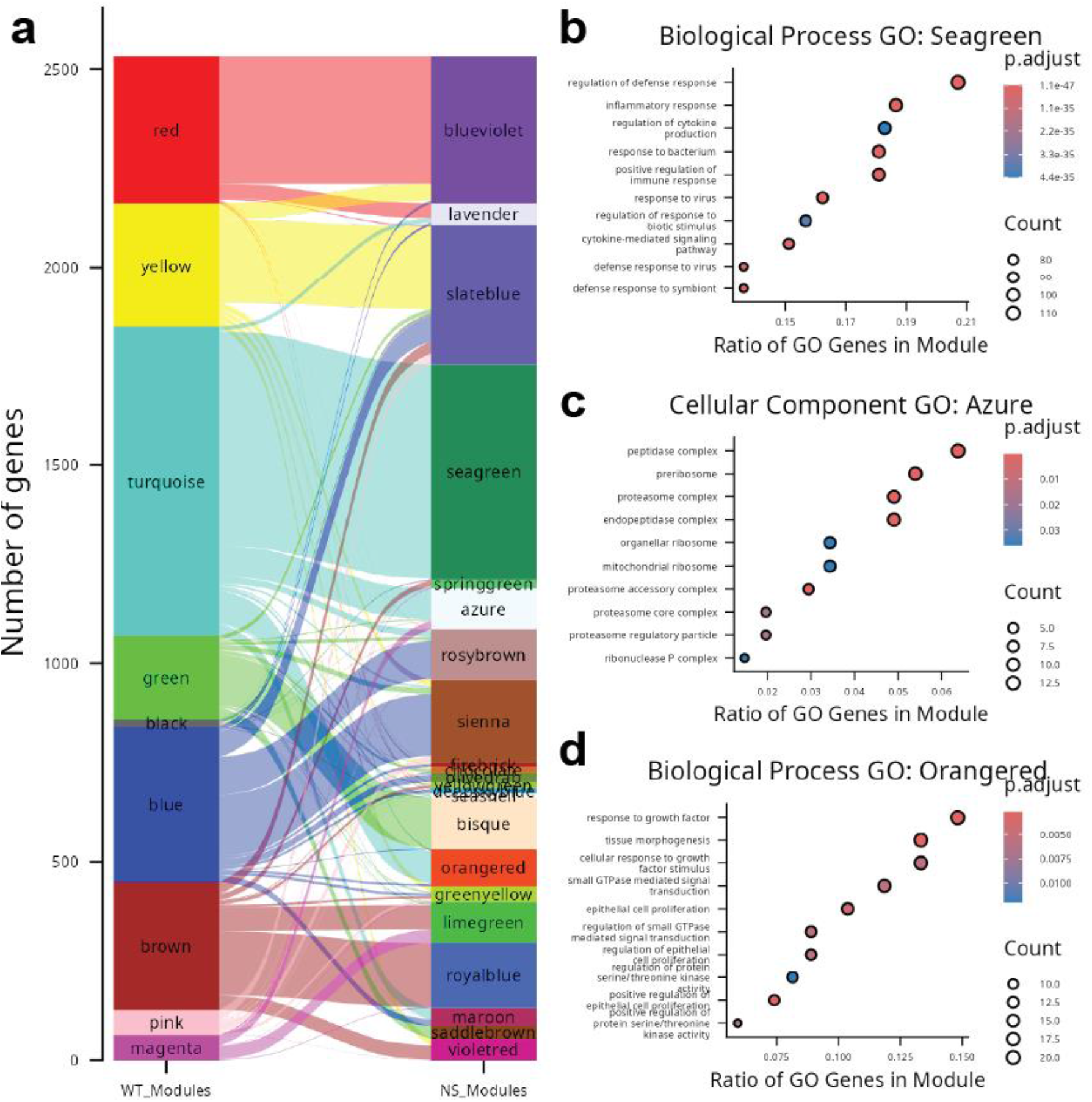
(a) Module membership of genes across hdWGCNAs of WT to WT+NS. The left column shows the genes and their identity in the WT the left column shows the same genes and their identity in the WGCNA performed across all backgrounds. (b,c,d) Gene Ontology Enrichment Analysis was performed on modules that share a majority of their genes with the WT PIC turquoise module. The dot plot shows the ten enriched terms with the lowest adjusted p value, represented by the dot’s color. The amount of genes associated with the particular term is represented by the dot’s size. The x axis represents the ratio of genes associated to the GO Biological Process or Cellular Compartment ontology term as a percent of the total genes belonging to the module.

### ZIKV NS genes modulate antiviral response submodules

Having characterized transcriptomic modules that represented the antiviral response using population level correlation across all conditions, we then looked to see whether ZIKV NS proteins were able to impact module activity within their respective sub-populations. To accomplish this, we performed a differential module expression score analysis, in which the mean expression score of each module is compared against PIC treated GFP transduction control (**Fig 4a**). Confirming the method, the canonical antiviral and proteasome modules were downregulated in unstimulated CTRL samples across all backgrounds.

**Figure 4.4.**
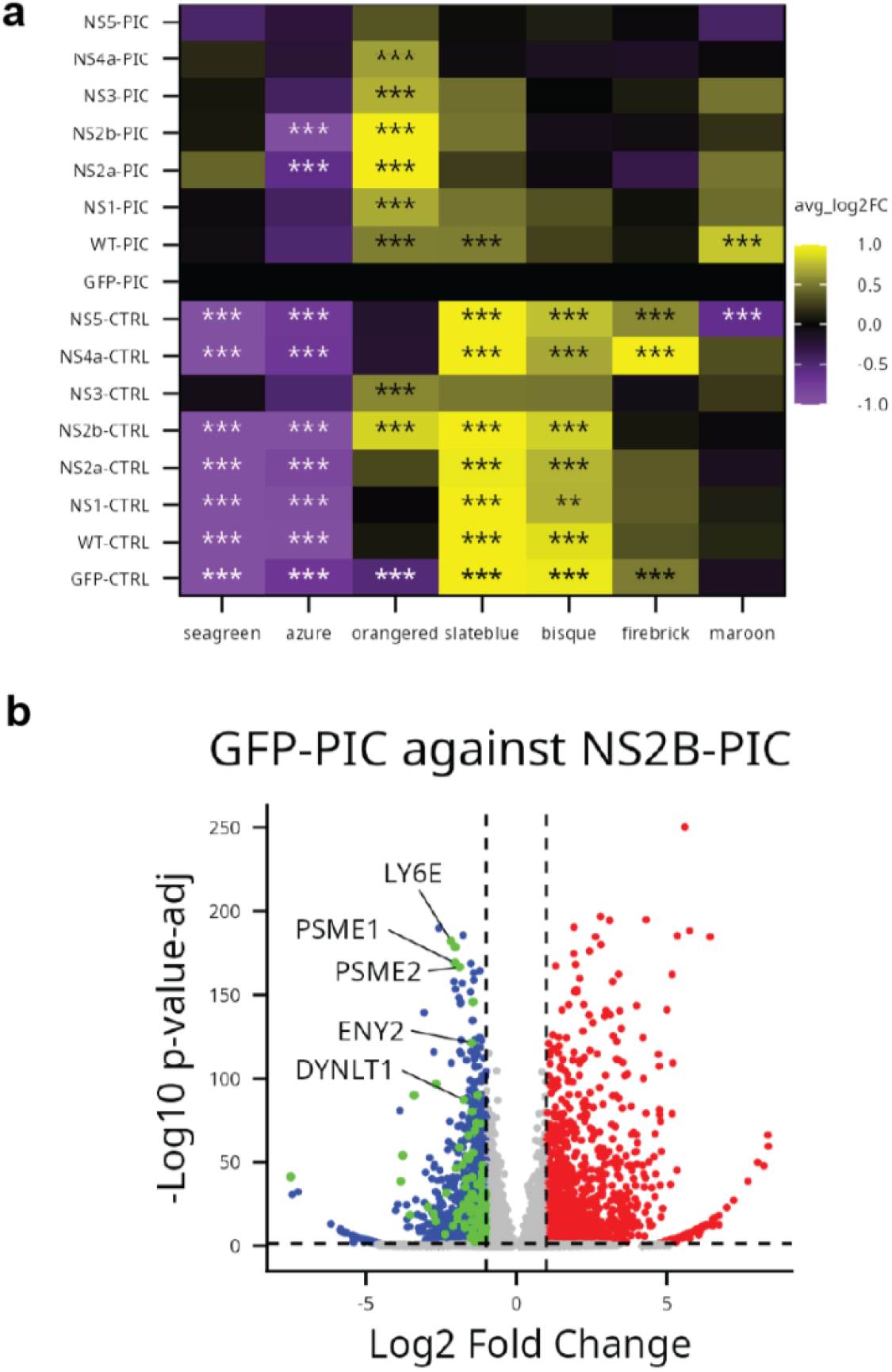
NS2B Reverses Proteasomal Activator Module Upregulation by PIC treatment. (a) Module expression scores were calculated and compared against expression scores in the GFP-PIC sample. Each row represents a sample and each column a module. 2 Log fold change is represented by each square’s color, and asterisk are superimposed over the square with a significant expression score difference, in respect to GFP-PIC. (b)Network graph portraying genes of the azure module as nodes and expression similarity scores as edges. Plotted edges represent the highest 2% of expression similarity scores. Genes differentially downregulated in NS2B-PIC rl=elative to GFP-PIC are colored red.

Notably, a significant downregulation in the expression score of the proteasome submodule was observed in the NS2b-PIC treated sample. Despite PIC stimulation, NS2b presence restores module expression score to levels similar to those of untreated samples, suggesting a strong modulation of module activity. Following up on GO terms we found proteasome subunit alpha type-3 (PSMA3) an integral protein to the 20S proteasome, and proteasome activator subunits 1 and 2 (PSME1 and PSME2), the immunoproteosome activation complex PA28, among the top ten module hub genes (Huber and Groll 2017).

Since module expression scores are calculated from average expression of all genes inside a module, it is possible that they can be influenced by large changes in expression of genes at the periphery of the module network as opposed to changes in hub genes more representative of the module. To address this we performed DE analysis comparing GFP-PIC to the NS2b-PIC sample and cross referenced the results with the hub genes of the proteasome module. Of the 90 genes belonging to the proteasome module found to be significantly downregulated in NS2b-PIC, 12 of them were considered hub genes, including the PA28 subunits PSME1 and PSME2 (**Fig 4b**).

## DISCUSSION

In this study we applied reductionist experimental design with a systems readout to dissect ZIKV bioactivity during an innate immune, antiviral response. Time restricted, induced expression of individual ZIKV NS genes allowed us to examine their effects individually and provided contrastive conditions that effectively split the monolithic antiviral response into smaller, biologically relevant functional units. HMC3 cells were the chosen model for their similarity to microglia, but the mechanisms identified were common to almost every cell type. While each NS protein had unique and measurable effects, NS2b presented a clear mechanism that fits known biology, but had not been previously reported. NS2B suppression of the immunoproteosome and MHCI display by downregulating expression of both PA28 cap elements, PSME1 and 2.

While single cell transcriptomics constitutes the richest available modality to capture systems level effects, microglial transcriptomic responses are highly dynamic and multidimensional, comprising a continuous co-expression landscape, rather than clear states for direct comparison (Dubbelaar et al., 2018; Paolicelli et al., 2022). Weighted Gene Co-expression Network Analysis (WGCNA) provided a systems framework to cluster co-varying genes into functional modules, capturing higher-order biological states beyond individual differential gene expression (Langfelder & Horvath, 2008; Morabito et al., 2023). By evaluating Module Eigengenes (MEs) and module preservation statistics across experimental conditions, network analysis can resolve subtle pathway discoordination, submodule splitting, or network disintegration caused by individual ZIKV non-structural proteins (Langfelder & Horvath, 2008; Morabito et al., 2023; Albertini et al., 2026). Integrating co-expression modules with transcription factor (TF) regulon and promoter motif enrichment analysis allows identification of core upstream master regulators that govern microglial antiviral submodules and are targeted by viral immunomodulators (Plaisier et al., 2016; Morabito et al., 2023.)

### Experimental approach: benefits and areas of opportunity

Combinatory perturbation of PIC stimulation and ZIKV NS1-5 protein transduction created biologically relevant variance that revealed novel mechanisms when parsed by hdWGCNA. ZIKV perturbation disturbed the coordination of an initially monolithic transcriptional response to PIC stimulation, allowing us to subdivide the antiviral response into three sub-modules. Studying individual module gene membership and activation across treatment conditions, we could infer an associated function and role in immune response. Following this approach, we identified three functions associated with the PIC response cell state. We also identified the extent to which these immune relevant modules were perturbed by individual ZIKV NS proteins and found a novel effect of PA28 inhibition by ZIKV NS2b.

While perturb-seq is an established method, applying hdWGCNA to perturbed scRNA-seq results in high sensitivity and resolution for identifying interference of cell functional responses by exogenous interventions (Adamson et al. 2016; Morabito et al. 2023). Such an experimental approach could be extended to further characterize immunomodulation of other viruses or molecules.

IFN-γ and IRF1, both upregulated in PIC response, increase the expression of immunoproteasome subunits PSMB8, PSMB9, and PSMB10, that replace the catalytic subunits β1,β2 and β5 found in constitutive proteasomes. The shift into immunoproteasome can lead to increased autoimmune activity and prolongs inflammatory activity (Miao et al. 2026). Along with changes to the proteasome catalytic subunits, IFN-γ also increases the expression of PSME1 and PSME2, subunits of the proteasome activator PA28. Binding of PA28 increases proteasome permissiveness to un-ubiquitinated proteins, especially oxidized proteins and small peptides, increasing their degradation (Wu et al. 2023). This degradation is specifically antiviral, both from the processing of viral proteins damaged by active oxidative immune processes and the increased oxidative stress from viral infection. The PA28 immunoproteasome also increases the ratio of degraded peptides that have a basic C-terminus, which have a higher affinity to MHC class 1 display proteins, promoting T-cell lead immune response and prolonging inflammatory conditions and increasing the probability of an adaptive immune response (Thomas et al. 2023).

Immunoproteasome dynamics after immune stimulation were recapitulated in our own data. All catalytic subunits of the immunoproteasome (PSMB8-10) were upregulated after PIC treatment and in all cases but one, PA28 expression was likewise upregulated after immune stimulation. Only NS2b caused a divergence in expression between the immunoproteasome and the PA28 subunits, PSME1 and PSME2. This NS2b, PA28 interaction is a novel finding and presents opportunities for further development in antiviral treatments or ZIKV bioengineering.

Proteasome interference and cooptation as a strategy for immune evasion has been widely observed in viruses (Wu et al. 2025). Viral inhibition of host PA28 formation specifically, has previously been described in the case of HIV’s p24 protein (McCarthy and Weinberg 2015). Viral PA28 inhibition leads to decreased antigen presentation in dendritic cells. Similar PSME1 and PSME2 inhibition by ZIKV, then, might impair the innate immune antiviral response by slowing down clearance of oxidized viral proteins during infection, as well as impairing the adaptive immune response by decreasing T-cell activation against ZIKV antigens.

Compounds for the stabilization of the PA28 activator are currently being developed to enhance T-cell activity against myeloma (Rana et al. 2024). If PA28 inhibition is found to be a significant strategy of ZIKV’s immune evasion, PA28 stabilizers could be repurposed as novel antivirals. Conversely, ZIKV immune evasion might be leveraged as a virally directed mechanism for antigen display inhibition in the treatment of autoreactive diseases. Like in the case of autoimmune diabetes, where peptide based MHC class-I inhibition has been seen to protect islet cell populations in mice (von Herrath et al. 1998).

In the brain, the immunoproteasome primarily occurs in astrocytes and microglial cells both capable antigen display, upon induction by TNF-α and IFN-γ (Puttaparthi and Elliott 2005). The immunoproteasome may either play a protective role or drive neuroinflammation with excessive activation causing cytokine release (Orre et al. 2013). In mouse models, immunoproteasome-deficient microglia engage and engulf tau protein aggregates, but fail to efficiently resolve internalized cargo and reducing cognitive function (Srikanth et al. 2026). However in other studies, inhibiting the immunoproteosome reduced the release of inflammatory factors and improved cognitive function (Yeo et al. 2019). Increased MHCI display is also inflammatory under the right circumstances, allowing increased invasion by CD8+ T cells (Seady et al. 2026). While a necessary part of immune function, chronic inflammation is an established driver of neurodegenerative pathologies with increased display as a hallmark of “inflammaging” and senescence (Kellogg et al. 2023).

For microglia specifically, MHCII display is not a homeostatic function, nor are microglia considered dendritic like cells, but with sufficient immune stimulus will activate MHCII display to interact with CD4+ and CD8+ T cells (Schetters et al. 2018). While partitioned away from MHCI mechanisms, both display systems are subject to cross talk and interact with T cells to shape systemic inflammation (Mantegazza et al. 2013).

### Limitations of single ZIKV protein transduction as model to study ZIKV immunomodulation

Our single collection of transcriptomic data at 24 hours after stimulation limits our ability to robustly capture early innate antiviral response. Recognition of PIC by TLR3 and RIG-I induces expression of pro-inflammatory genes 3-5 hrs after stimulation. The second large transcriptomic event in innate antiviral response occurs at around 24 hours post stimulation, when IFN induced expression begins (Valdés-López et al. 2025). We elected for a 24 hr time point as it is the earliest likely moment to capture transcriptomic evidence of both early and late responses. The ∼20 hr gap between the start of TLR3-RIG-I signaling and collection is twice as long as the median half life of mRNA (Yang et al. 2003). This meant a loss of resolution on early transcriptomic effects of ZIKV NS1-5 interference as mRNA degradation rates influenced our readout. Notably, no IFN transcripts were detected, only response genes. Collection at earlier timepoints would be necessary to resolve this issue, at the expense of capturing later responses, but could reveal further bioactivity.

While interference during the early stage could have measurable effects on downstream IFN production and signaling, we did not observe ZIKV NS presence significantly alter the majority of IFN dependent expression captured in the seagreen module. This might be due to the fact that early activation has redundant pathways, while individual ZIKV NS proteins only target one or two at a time. Presence of the entire ZIKV genome might be necessary to significantly impede transition of early to late innate response.

The function of each module inferred from GO term association and hub gene membership, but validated by secondary assays. Assertions of functional links could be strengthened by using transcription factor association pipelines to predict what transcription factors may be coordinating module expression.

Finding such coordination would show that these genes are not simply covariant by random chance but due to a specific transcriptional factor’s activity. Implicating a real biological cause and a testable mechanism.

To mechanistically link module coordination and activity to cellular function non-transcriptomic measurements are needed. For example, an approach in validating the apoptosis module’s association to actin cytoskeleton maintenance could be to perform spatial transcriptomics alongside immunocytochemistry or cell morphology analysis. In this way the functional association could be tested by correlating transcriptomic changes of apoptosis hub genes to cell roundedness or total actin content.

## METHODS

### Tissue Culture

HMC3 (CRL-3304) and HEK293 (CRL-1573) cells were grown on DMEM 4.5g/L glucose with Glutagro (10-101-CV) 10% FBS and 1% Penicillin/Streptomycin in an incubator set at 37C and 5% CO2.

### Recombinant Plasmid Construction

ZIKV (strain: ZKC2) non-structural (NS) genes were synthesized and isolated by PCR. The linear fragments were designed to have the exact amino acid sequence of its protein cognate, following ZIKV polyprotein processing seen after infection. This meant including membrane binding domains that might technically lie in the preceding NS gene. The transmembrane domains align the viral proteins on either end of the endoplasmic reticulum (ER) membrane. The isolated fragments were then cloned into the lentiviral backbone, pLenti, using the NEB Hi-Fi DNA Assembly kit. Downstream of the ZIKV NS gene, an IRES region followed by the enhanced green fluorescent protein (EGFP) were included; creating a polycistronic cassette of ZIKV NS gene and a fluorescent reporter. The cassette was placed under the expression of the tetracycline inducible promoter TRE3G. To enable tetracycline induction, we cloned the active reverse tetracycline-controlled transactivator (rtTA) under expression of the constitutive promoter PGK.

### Transduction and Selection

Each ZSC plasmid, alongside the lentiviral helper plasmids pMD2G and psPAX2, were transfected in equimolar concentration into 2 6-wells of HEK293 cells at a confluency of ∼60% using Lipofectamine 3000 (cat:L3000001) per supplier instructions. 72 hours post transfection, media was collected, pooled and filtered through a 0.45 µm syringe filter. 4ug/ml of polybrene was added to the viral media and then diluted 1:5 using fresh complete DMEM. To transduce our ZIKV NS lines, 2 mls of diluted viral media was added onto HMC3 cells plated in a 6 well plate at ∼40% confluency.

### Sorting of Transduced Cells

48 hrs after transduction, expression of the indubile ZSC NS/EGFP polycistron was induced by addition of 10μg/ml doxycycline. 48 hours after induction, cells were separated using a BD FACSymphony S6 cell sorter, with EGFP fluorescence as the selection marker. Sorted cells were then pooled into 96 wells and grown back to confluence. Having reached confluence, the undiluted cells were passaged into a 48 well plate. This process of up-passaging was repeated until we reached a confluent transduced culture in a T75 flask. At which point cells were then prepared for induction and scRNA collection. Frozen stocks were created using unused cells.

### scRNA Collection

Cells were passaged into a 6 well plate at 50% confluence. A day after plating, cells were incubated with 10μg/ml doxycycline for 24 hrs.Media was then replaced with DMEM containing 10μg/ml doxycycline as well as one of three immune stimulation treatments, 50 ul of PBS for control, 100μg/ml LPS or 10μg/ml PIC. 24 hrs after immune stimulation cells were collected and prepped for scRNA seq using the Illumina Single Cell 3′ RNA Prep, T2 kit (cat: 20135689). Transcriptomic data was collected from cells at passage number 15-20. Samples were sequenced in a paired end 150 bp sequencing format on a single NovaSeq XPlus 25B Lane.

### Sequence alignment

To permit quantification of transgenic transcripts during alignment, a custom transgenic reference chromosome was appended to the human reference assembly (Ensembl hg38 v115). The custom reference chromosome consisted of the IRES-EGFP region of the polycistronic cassette, spanning 1489 bp. The combined genomic reference assembly was then used to create reference files for single cell transcriptomic alignment using the DRAGEN tool, Reference Builder (v. 4.4.6). Sequence alignment, cell barcoding, and unique molecular identifier (UMI) deduplication was executed using the DRAGEN Single Cell RNA application.

### Sample QC and Filtering

Aligned fastqc files were converted into Seurat objects using the R Seurat package (5.5.1). Quality control checks were performed to remove any dying cells or doublets. Mitochondrial gene percentages were used as a proxy of cell quality. We assessed plots of mitochondrial gene percentage against the number of RNA molecules in a cell by cell basis to set threshold limits on mitochondrial gene percentages. Thresholds varied across samples but were set at a lower limit of 0.2-0.3% to an upper limit of 2-3%. To remove suspected doublets, cells with number of features and number of RNA molecules well above the mean were manually thresholded out.

Samples were subset to only include cells with more than 10,000 RNA molecules, so as to produce samples with similar levels of read depth and decrease sparsity that would skew measures of covariance. Normalization, data scaling, and identification of most variable features was performed using the Seurat function SCTransform. PCA and UMAP were performed using the “SCT” assay. Data showed overlap by background and segregation by treatment (**Fig S3**).

### Differential Expression Analysis and Gene Ontology enrichment

Differentially expressed genes were identified using the Seurat function FindMarkers(). Significant differential expression was defined as any genes that experienced a 2 fold change and received an adjusted p value below 0.05.

Genes lists set for gene ontology (GO) enrichment were converted to ENTREZ ids, conversions were checked to ensure no gene list ever saw a loss of genes above 5%. GO enrichment was performed using the package ClusterProfiler (4.10.1) and the Bioconductor human ontology annotation database org.HS.eg.db. A q-value cutoff of 0.05 was used to determine significant enrichment. Hypergeometric testing was performed using the base R phyper function. Any resulting p values below the minimum representable value are shown as p<2.2^-308.^.

### Metacell construction and hdWGCNA

Before hdWGCNA processing, subset data was further subset by eliminating all genes from expression data that did not appear in at least 5% of cells. To combat sparsity inherent in single cell transcriptomic datasets and improve signal-to-noise ratio, k-nearest neighbor cell pooling was utilized to construct metacells within biological groups. Metacells were constructed using the hdWGCNA function MetacellByGroups,with the following parameters: k.parameter=25, max_shared=5. Metacells construction was made by grouping by samples.

High Dimensionality Weighted Gene Correlation Network Analysis (hdWGCNA) was performed on constructed metacells, using the function ConstructNetwork from the hdWGCNA R package (0.4.12). The parameters used for a signed network construction are as follows: merge_height=0.15, deep_split=3, mod_size=50, net.type=”signed”. The soft power threshold was selected at the point at which fit to a scale free model exceeds 0.8, soft power threshold crawl was performed using the function TestSoftPowers.

### Module Preservation Test

Modules discovered in WT hdWGCNA were projected onto all other samples grouped by genetic background. Module preservation tests were performed using the hdWGCNA function Module Preservation() for 150 permutations. Density tests measure whether to what extent query modules remain covariant to one another in the tested dataset. Separability tests measure to what extent the modules from the query dataset remain distinct in the tested dataset. Summary scores combine both measures to represent overall module preservation. Z scores are produced by performing measurements of module preservation in a tested dataset using randomly assigned genes and genes belonging to the queried modules. The difference in distribution is used to calculate the probability with which the module genes perform better on these measures than any random assortment of genes (Langfelder et al. 2011).

## SUPPLEMENTAL MATERIALS

**Supplemental Figure 4.1.**
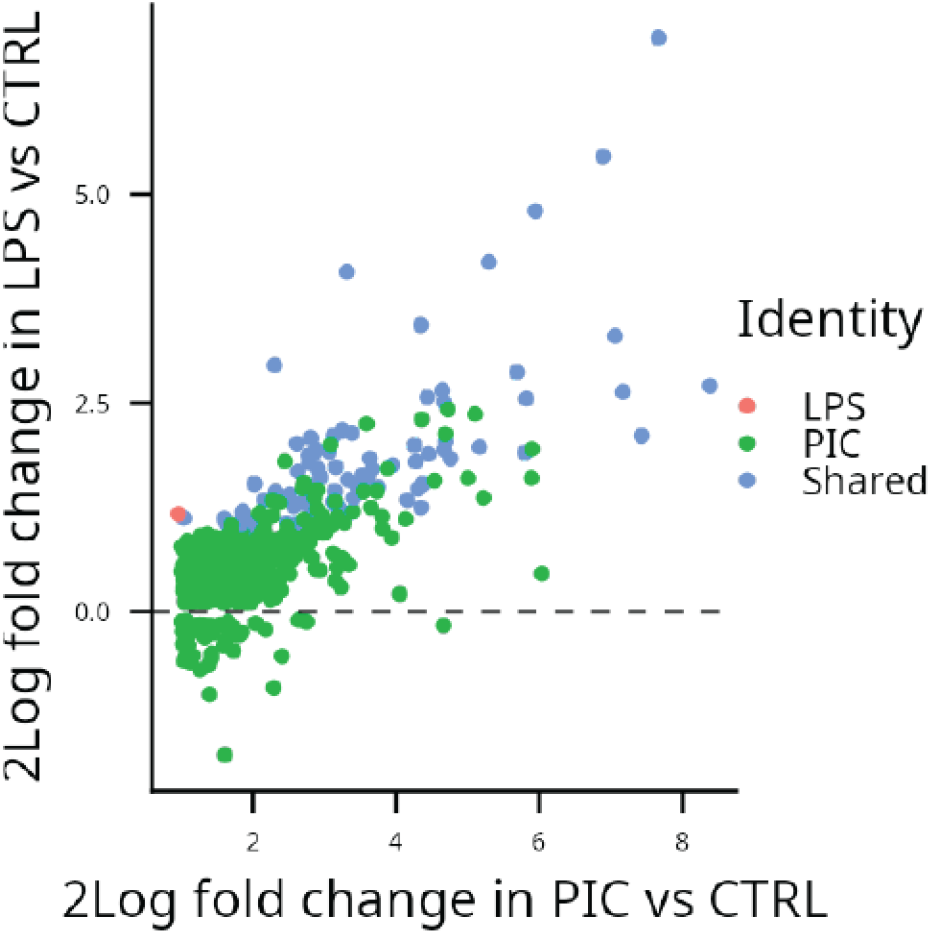
Overlap of PIC and LPS differentially upregulated genes. Dot plat showing all differentially upregulated genes in WT-CTRL vs WT-PIC and WT-LPS. Genes that are only differentially upregulated in LPS treatment are colored, PIC in green and genes that are differentially upregulated in both DE analysis are colored blue

**Supplemental Figure 4.2.**
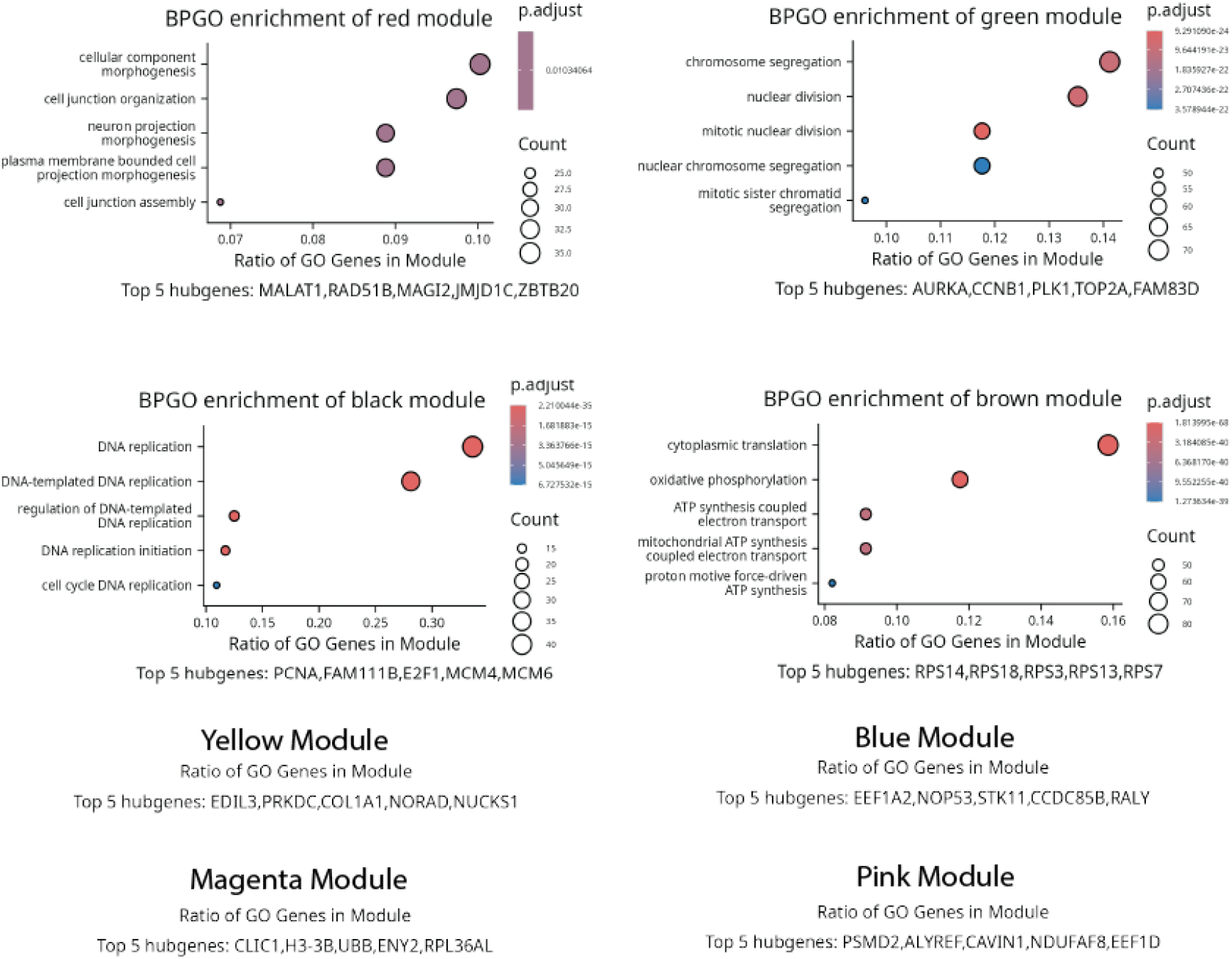
Enriched GOs and hub genes of WT hdWGCNA modules. Hub genes are shown at the bottom of each plot. No plot is shown for modules that did not have any significantly associated GO terms.

**Supplemental Figure 4.3.**
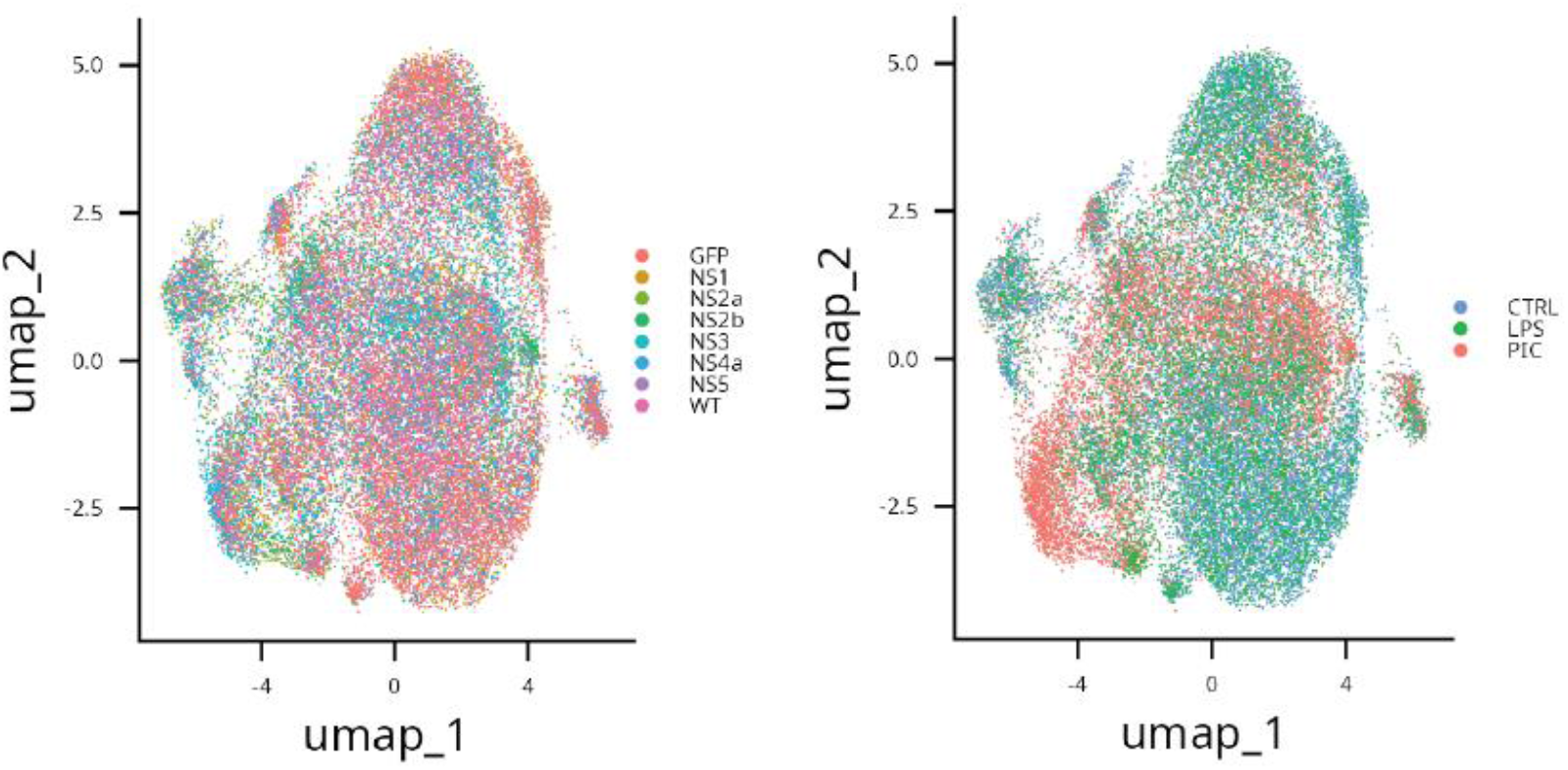
UMAP of samples grouped by treatment and genetic background.

**Supplemental Figure 4.3.**
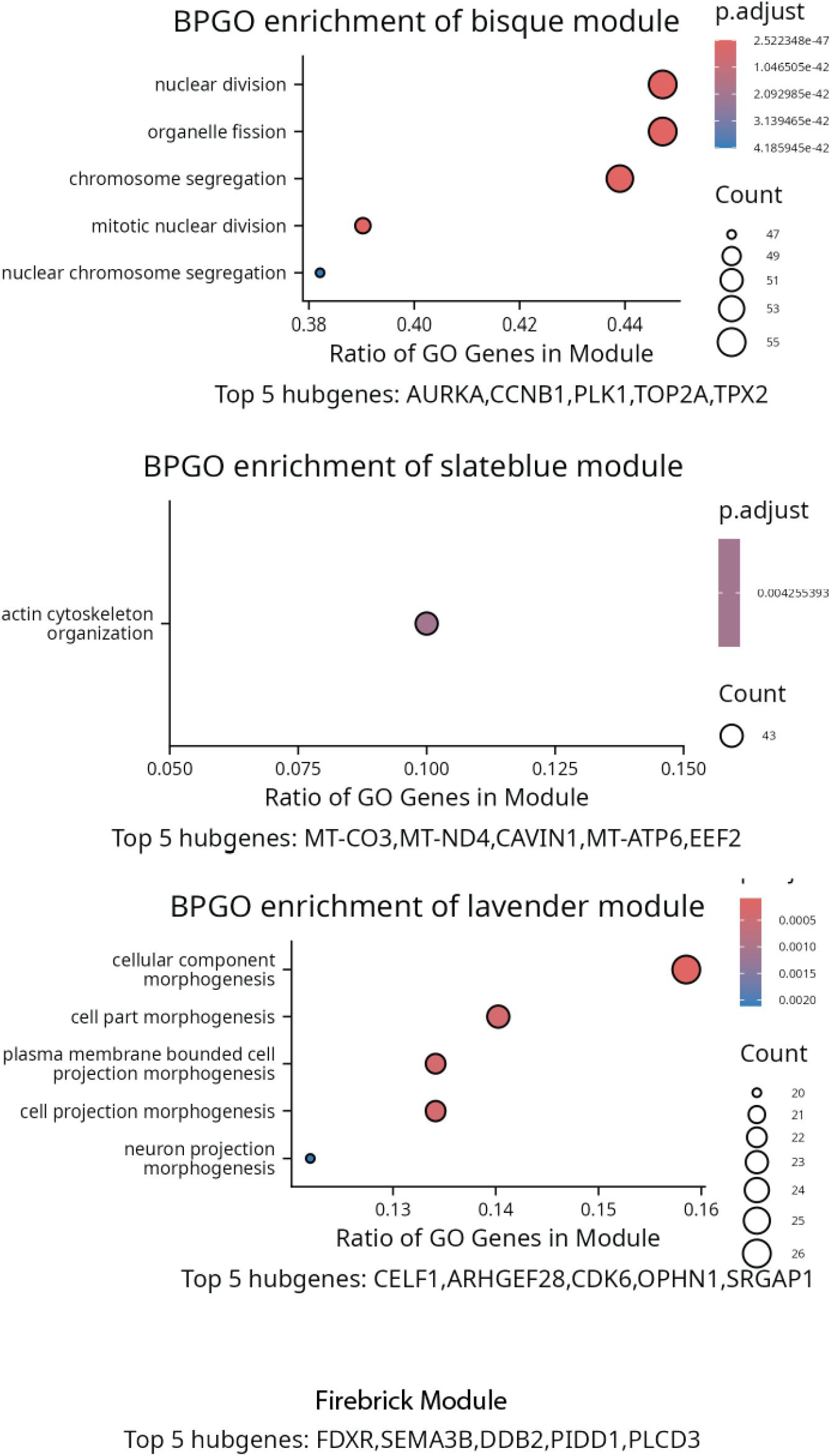
Enriched GOs and Hub Genes of Homeostatic Submodules. Hub genes are shown at the bottom of each plot. No plot is shown for modules that did not have any significantly associated GO terms.

## REFERENCES

Adamson B, Norman TM, Jost M, Cho MY, Nuñez JK, Chen Y, Villalta JE, Gilbert LA, Horlbeck MA, Hein MY, et al. 2016. A Multiplexed Single-Cell CRISPR Screening Platform Enables Systematic Dissection of the Unfolded Protein Response. Cell. 167(7):1867–1882.e21. 10.1016/j.cell.2016.11.048

Ahmad L, Zhang S-Y, Casanova J-L, Sancho-Shimizu V. 2016. Human TBK1: A Gatekeeper of Neuroinflammation. Trends Mol Med. 22(6):511–527. 10.1016/j.molmed.2016.04.006

Fan X, Diao W, Wang H, Yin X, Qian W. 2025. Interferon Regulatory Factors as a Potential Therapeutic Target for Neuroinflammation: A Focus on Alzheimer’s Disease. Int J Mol Sci. 26(7):2906. 10.3390/ijms26072906

He Y, Taylor N, Yao X, Bhattacharya A. 2021. Mouse primary microglia respond differently to LPS and poly(I:C) in vitro. Sci Rep. 11:10447. 10.1038/s41598-021-89777-1

von Herrath MG, Coon B, Lewicki H, Mazarguil H, Gairin JE, Oldstone MBA. 1998. In Vivo Treatment with a MHC Class I-Restricted Blocking Peptide Can Prevent Virus-Induced Autoimmune Diabetes. J Immunol. 161(9):5087–5096. 10.4049/jimmunol.161.9.5087

Huber EM, Groll M. 2017. The Mammalian Proteasome Activator PA28 Forms an Asymmetric α4β3 Complex. Structure. 25(10):1473–1480.e3. 10.1016/j.str.2017.07.013

Kellogg CM, Pham K, Machalinski AH, Porter HL, Blankenship HE, Tooley KB, Stout MB, Rice HC, Sharpe AL, Beckstead MJ, et al. 2023. Microglial MHC-I induction with aging and Alzheimer’s is conserved in mouse models and humans. GeroScience. 45(5):3019–3043. 10.1007/s11357-023-00859-6

Langfelder P, Luo R, Oldham MC, Horvath S. 2011. Is my network module preserved and reproducible? PLoS Comput Biol. 7(1):e1001057. 10.1371/journal.pcbi.1001057

Lawrence JH, Sherer ML, Tavlarides-Hontz P, Parcells MS, Schwarz JM. 2019. An investigation into the immune response of cultured neural rat cells following Zika virus infection. J Neuroimmunol. 332:73–77. 10.1016/j.jneuroim.2019.04.002

Mantegazza AR, Magalhaes JG, Amigorena S, Marks MS. 2013. Presentation of Phagocytosed Antigens by MHC Class I and II. Traffic. 14(2):135–152. 10.1111/tra.12026

McCarthy MK, Weinberg JB. 2015. The immunoproteasome and viral infection: a complex regulator of inflammation. Front Microbiol. 6:21. 10.3389/fmicb.2015.00021

Meertens L, Labeau A, Dejarnac O, Cipriani S, Sinigaglia L, Bonnet-Madin L, Le Charpentier T, Hafirassou ML, Zamborlini A, Cao-Lormeau V-M, et al. 2017. Axl Mediates ZIKA Virus Entry in Human Glial Cells and Modulates Innate Immune Responses. Cell Rep. 18(2):324–333. 10.1016/j.celrep.2016.12.045

Miao J, Shaheen N, Zhao Y, Zhao J. 2026. Emerging role of the immunoproteasome as a druggable target in lung inflammatory diseases. Respir Res. 27(1):102. 10.1186/s12931-026-03524-6

Morabito S, Reese F, Rahimzadeh N, Miyoshi E, Swarup V. 2023. hdWGCNA identifies co-expression networks in high-dimensional transcriptomics data. Cell Rep Methods. 3(6):100498. 10.1016/j.crmeth.2023.100498

Orre M, Kamphuis W, Dooves S, Kooijman L, Chan ET, Kirk CJ, Dimayuga Smith V, Koot S, Mamber C, Jansen AH, et al. 2013. Reactive glia show increased immunoproteasome activity in Alzheimer’s disease. Brain. 136(Pt 5):1415–1431. 10.1093/brain/awt083

Puttaparthi K, Elliott JL. 2005. Non-neuronal induction of immunoproteasome subunits in an ALS model: possible mediation by cytokines. Exp Neurol. 196(2):441–451. 10.1016/j.expneurol.2005.08.027

Rana PS, Ignatz-Hoover JJ, Guo C, Mosley AL, Malek E, Federov Y, Adams DJ, Driscoll JJ. 2024. Immunoproteasome Activation Expands the MHC Class I Immunopeptidome, Unmasks Neoantigens, and Enhances T-cell Anti-Myeloma Activity. Mol Cancer Ther. 23(12):1743–1760. 10.1158/1535-7163.MCT-23-0931

Schetters STT, Gomez-Nicola D, Garcia-Vallejo JJ, Van Kooyk Y. 2018. Neuroinflammation: Microglia and T Cells Get Ready to Tango. Front Immunol [Internet]. [accessed 2026 Oct 24] 8. 10.3389/fimmu.2017.01905

Seady M, Maynes MA, Thelwell JS, Jin F, Hansen MJ, Jensen HE, Witter RK, Owens CA, Hassani A, Lewis CL, et al. 2026. TMEM119+ microglia MHC class I restricted antigen presentation impacts CD8 T cell memory, effector status, and blood-brain barrier disruption during neurotropic virus infection. bioRxiv.:2026.05.08.722741. 10.64898/2026.05.08.722741

Srikanth M, Jiang S, Wellman SM, Sarkar S, Lorman DE, Lantin T, Runyan AM, Kumar M, Sydney E, Figueroa HY, et al. 2026. Immunoproteasome Deficiency Impairs Microglial Clearance and Worsens Tau and Amyloid Pathology [Internet]. [accessed 2026 Nov 7]:2026.07.14.738427. 10.64898/2026.07.14.738427

Thomas T, Salcedo-Tacuma D, Smith DM. 2023. Structure, Function, and Allosteric Regulation of the 20S Proteasome by the 11S/PA28 Family of Proteasome Activators. Biomolecules. 13(9):1326. 10.3390/biom13091326

Valdés-López JF, Tamayo-Molina YS, Fernandez GJ, Hernández-Sarmiento LJ, Velilla PA, Urcuqui-Inchima S. 2025. Early events in dengue virus infection inducing cytokine storm: The dynamic interplay of pattern-recognition receptors, inflammasome activation, and biphasic NF-κB and STAT1-dependent inflammatory responses in human mononuclear phagocytes. PLoS Negl Trop Dis. 19(9):e0013366. 10.1371/journal.pntd.0013366

Wu D, Wang Y, Zhou Y, Gao H, Zhao B. 2023. Inhibition of the Proteasome Regulator PA28 Aggravates Oxidized Protein Overload in the Diabetic Rat Brain. Cell Mol Neurobiol. 43(6):2857–2869. 10.1007/s10571-023-01322-y

Wu Y, Sun Z, Xia L, Tian P, Jiao L, Li Y, Wei Z, Wang X, Li X, Zhang G. 2025. MHC-I pathway disruption by viruses: insights into immune evasion and vaccine design for animals. Front Immunol [Internet]. [accessed 2026 Oct 24] 16. 10.3389/fimmu.2025.1540159

Yang E, van Nimwegen E, Zavolan M, Rajewsky N, Schroeder M, Magnasco M, Darnell JE. 2003. Decay Rates of Human mRNAs: Correlation With Functional Characteristics and Sequence Attributes. Genome Res. 13(8):1863–1872. 10.1101/gr.1272403

Yeo IJ, Lee MJ, Baek A, Miller Z, Bhattarai D, Baek YM, Jeong HJ, Kim YK, Kim D-E, Hong JT, Kim KB. 2019. A dual inhibitor of the proteasome catalytic subunits LMP2 and Y attenuates disease progression in mouse models of Alzheimer’s disease. Sci Rep. 9(1):18393. 10.1038/s41598-019-54846-z

Zhang R-Z, Li L, Ma C, Dou T-T, Wei Y-T, Yan X-K. 2025. Advances in the Understanding of Mitochondrial Inflammatory Regulation in the Pathogenesis of Alzheimer’s Disease. J Inflamm Res. 18:14475–14491. 10.2147/JIR.S557000

